# The BRadykinesia Akinesia INcoordination (BRAIN) tap test: capturing the sequence effect

**DOI:** 10.1101/453852

**Authors:** Hasan Hasan, Maggie Burrows, Dilan S. Athauda, Bruce Hellman, Ben James, Tom Warner, Thomas Foltynie, Gavin Giovannoni, Andrew J. Lees, Alastair J. Noyce

## Abstract

**Background:** The BRAIN tap test is an online keyboard tapping task that has been previously validated to assess upper limb motor function in Parkinson’s disease (PD).

**Objectives:** To develop a new parameter which detects a sequence effect and to reliably distinguish between PD patients ‘on’ and ‘off’ medication. Alongside, we sought to validate a mobile version of the test for use on smartphones and tablet devices.

**Methods:** BRAIN test scores in 61 patients with PD and 93 healthy controls were compared. A range of established parameters captured speed and accuracy of alternate taps. The new VS (Velocity Score) recorded the inter-tap speed. Decrement in the VS was used as a marker for the sequence effect. In the validation phase, 19 PD patients and 19 controls were tested using multiple types of hardware platforms including smart devices.

**Results:** Quantified slopes from the VS demonstrated bradykinesia (sequence effect) in PD patients (slope cut-off −0.002) with sensitivity of 58% and specificity of 81% (discovery phase of the study) and sensitivity of 65% and specificity of 88% (validation phase). All BRAIN test parameters differentiated between ‘on’ medication and ‘off’ medication states in PD. Most BRAIN tap test parameters had high test-retest reliability values (ICC>0.75). Differentiation between PD patients and controls was possible on all hardware versions of the test.

**Conclusion:** The BRAIN tap test is a simple, user-friendly and free-to-use tool for assessment of upper limb motor dysfunction in PD, which now includes a measure of bradykinesia.

## Introduction

The use of technology to complement clinical assessment of Parkinson’s disease (PD) is growing rapidly. Rating scales are valuable for clinical practice and research, but are prone to both inter- and intra-rater variability [1,2]. In order to obviate these shortcomings, a range of technologies measuring bradykinesia in PD have been developed [3–12].

The BRAIN (BRadykinesia Akinesia INcoordination) test is a freely-available online keyboard finger tapping test that is based on the alternate finger tapping task [13,14]. It has previously been shown to differentiate PD patients from healthy controls and has been used for longitudinal monitoring of motor function in the PREDICT-PD study, a large cohort of healthy older individuals stratified for future risk of PD [15].

In this work we developed a new parameter to quantify an aspect of bradykinesia (which is defined as “*slowness of initiation of voluntary movement with progressive reduction in speed and amplitude of repetitive actions*”) and known in motor physiology as the sequence effect [16]. We then used it to determine if it could reliably distinguish between patients with PD who were ‘on’ and ‘off’ dopaminergic medication. Finally, we validated it in a separate patient and control group and introduced the test to ‘smart’ devices.

## Methods

### Participants

For the first two experiments, we assessed 61 patients (61.3±8.2 years) with mild-moderate stage PD (Hoehn and Yahr <2.5) who were enrolled in the Exenatide-PD trial at the National Hospital, Queen Square, London. Inclusion and exclusion criteria for this trial have previously been published [9]. Retrospective data from 93 healthy age-matched controls (60.4±10.7 years) were used for comparison [15].

For the third experiment, 20 patients with PD (66.3±6.6 years) were recruited from a movement disorders clinic at the National Hospital, Queen Square. 20 healthy spouses (67.4±9.0 years) of the recruited patients acted as controls.

The studies were approved by the Brent NHS Research Ethics Committee, London (13/LO/1536) and the Queen Square Ethics Committee, London (09/HO716/48). Informed consent was obtained from all participants.

### Experimental procedure

To assess participants in the ‘off state’, patients were instructed to stop their medications for 12-36 hours prior to the study visit. Part III of the MDS-UPDRS was assessed alongside performance on the BRAIN tap test. Patients then took their regular medication. Measurements were repeated in the ‘on state’. The same neurologist (D.A.) performed all of the clinical ratings.

The BRAIN test experimental task has been described previously [14], and further information can be found here in the supplementary material (Pages S1-S2). Briefly, users are instructed to strike the ‘S’ and ‘;’ keys on a standard computer keyboard, alternately using one index finger, as fast and as accurately as possible, for 30 seconds.

Parameters generated from the test include KS (kinesia score – number of alternate taps in 30s), AT (akinesia time – mean dwell time on keys in milliseconds), IS (incoordination score – a measure of rhythm given by the variance in the travelling times between key presses) and DS (dysmetria score – a measure of the average accuracy of key strikes where the central key scores 1, adjacent keys are 2 and all other keys are 3).

In the third experiment, a smart device version of the test (“TapPD” developed by uMotif Limited for Apple iPhone and iPad devices) was used alongside the keyboard test [17][18]. Participants used their index finger to alternately tap two target areas on the screen as fast and as accurately as possible for a period of 30 seconds. The application captured the same measurements as the BRAIN test, but the dysmetria score was engineered to incorporate additional capabilities of smart devices. Accuracy of each tap within a hit area was calculated as a decimal, with 0 being at the centre of the target (perfect accuracy) and 1 being at the maximum edges of the hit area. DS1 was calculated as the average accuracy during the test. Screenshots of the “TapPD” interface can be viewed in the supplementary material (Figure S1).

We developed a new parameter, the Velocity Score (VS) by measuring the inter-tap velocity throughout the duration of the test. To look for a sequence effect in patients with PD, the percentage change in velocity with respect to the initial velocity between the first two key taps was computed and plotted as a time series graph. Slopes of acceleration/deceleration in the time-series graphs were compared between PD patients in the ‘off state’ and healthy controls (see Figure 1). The steeper the slope, the faster the rate in increase/decline of velocity over time. For simplicity, a linear trend line was used for slope quantification. Alternate approaches for calculating a sequence effect, using the number of consecutive decrements in dwell and travelling times (for example, three), were explored and are shown in the supplementary material (Pages S2-3, Figure S2).

**Figure 1.**
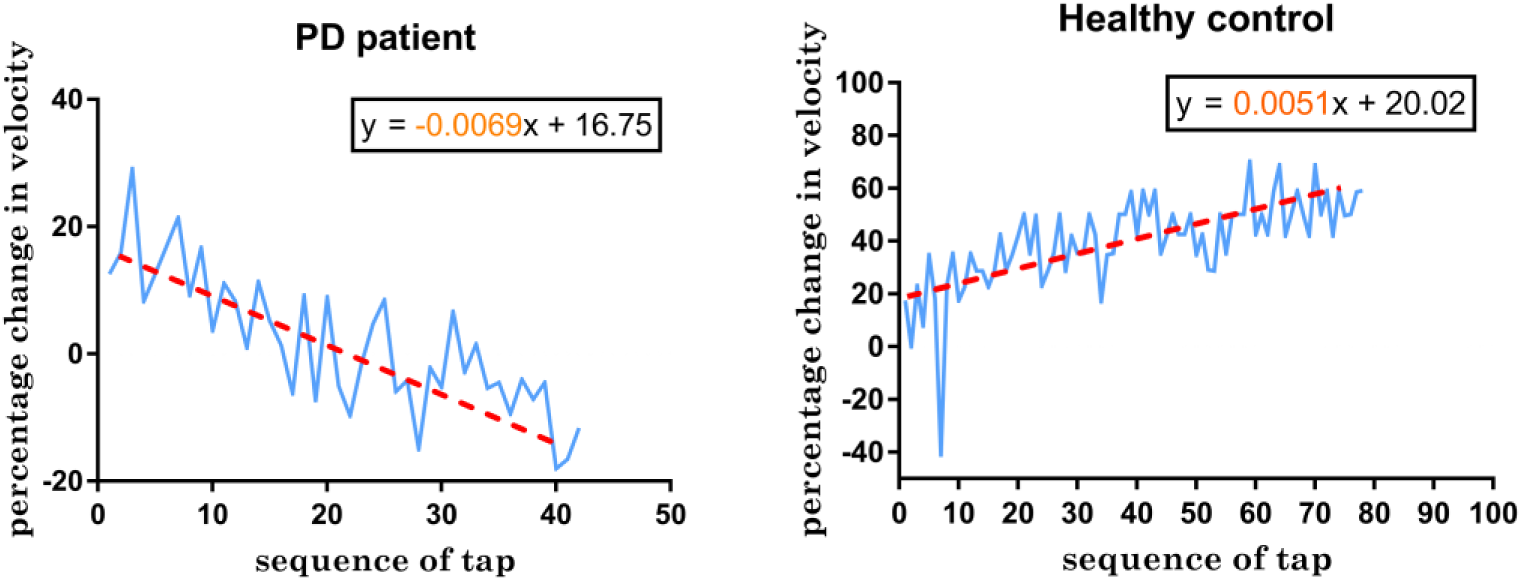
Time series analysis of change in velocity for the duration of the test compared to the initial velocity in a PD patient (left) and healthy control (right). The slope was derived from the regression equation of the linear trendline

In the third experiment, two trials of the BRAIN test were conducted for each hand on the computer keyboard, smartphone and tablet device.

### Statistical methods

Analyses were performed using GraphPad Prism statistical software (version 7.0) and IBM SPSS version 24 (SPSS Inc, Chicago, IL). A p value of ≤ 0.05 was used as cut-off for determining significance. To reduce the type I error resulting from multiple subgroup analyses, a false discovery rate (FDR) control for p values was used [19]. BRAIN tap parameters were correlated with total motor MDS-UPDRS part III score and sub-scores (rigidity, finger tapping, pronation-supination and hand movements) for the ‘on’ and ‘off’ states. Pearson’s correlation coefficient was used for normally distributed variables and Spearman’s rank correlation coefficient for non-parametric correlation. Normality was checked using the D’Agostino Pearson normality test. ROC (Receiver Operating Characteristic) curves were used to differentiate PD patients’ most affected side and controls’ least well performing score on the BRAIN tap test. Additionally, Wilcoxon matched pairs signed rank test was used to differentiate between PD patients’ scores ‘on’ and off’ medication. A Chi-square test for binary outcome variables was used to compare the epoch-analyses for the sequence effect between PD patients and controls. The Kruskal-Wallis test was used to compare the slopes the three groups (PD ‘on’, PD ‘off’, and controls). Fisher’s exact test was used to compare decrement in slopes between PD and controls.

In the third experiment, Intra-Class Correlation coefficient (ICC) for test-retest reliability was calculated based on single score, absolute-agreement, two-way mixed effects. Interpretation of ICC was conducted in accordance to the recommendations by Koo and Li - *<0.5* (poor reliability), *0.5-0.75* (moderate reliability), *0.75-0.9* (good reliability) and *>0.90* (excellent reliability)[20]. The Mann-Whitney U test was used to compare the distribution of test results between the two groups’ performance on the three platforms (keyboard, iPad and iPhone).

## Results

The demographic information for the participants is summarised in the supplementary material (see Table S1). One PD patient was excluded from MDS-UPDRS (total and motor sub-score) correlation with BRAIN tap scores in the second experiment due to incomplete data. In the third experiment, one PD patient and one control were excluded for technical reasons. Sensitivity and specificity cut-offs for test parameters are summarised in supplementary material (Table S2).

### Quantifying bradykinesia using slopes

In the discovery phase of the study, PD patients differed from controls (Fisher’s exact test p≤0.001) by showing decrements in the slope of velocity-time graphs (Table 1). Sensitivity and specificity were 58% and 81% respectively for a slope cut-off of −0.002, which is equivalent to a 1% decrement in velocity for every 5 alternate finger taps. Similarly, in the validation phase of the study, PD patients differed from controls (p = 0.004), with a sensitivity and specificity of 65% and 88% respectively using the same cutoff. A box and whiskers plot comparing slopes in PD and controls is shown in Figure 2. A slope of −0.002 in velocity-time graphs also corresponded to the average decrement in PD patients that had bradykinesia on clinical examination.

**Table 1.** Combined 2×2 contingency tables showing number of PD patients and controls with decrement and no decrement in velocity in experiment 1 and 3.

| Experiment 1<br>(discovery) | Decrement in<br>velocity | No decrement in<br>velocity | Total | P value |
| --- | --- | --- | --- | --- |
| Parkinson's disease<br>(most affected side) | 35 (58%) | 25 (42%) | 60 | <0.001 |
| Controls<br>(non-dominant side) | 11 (19%) | 47 (81%) | 58 |  |
| Experiment 3<br>(validation) | Decrement in<br>velocity | No decrement in<br>velocity | Total | P value |
| Parkinson's disease<br>(most affected side) | 11 (65%) | 6 (35%) | 17 | 0.0039 |
| Controls<br>(non-dominant side) | 2 (12%) | 15 (88%) | 17 |  |

**Figure 2.**
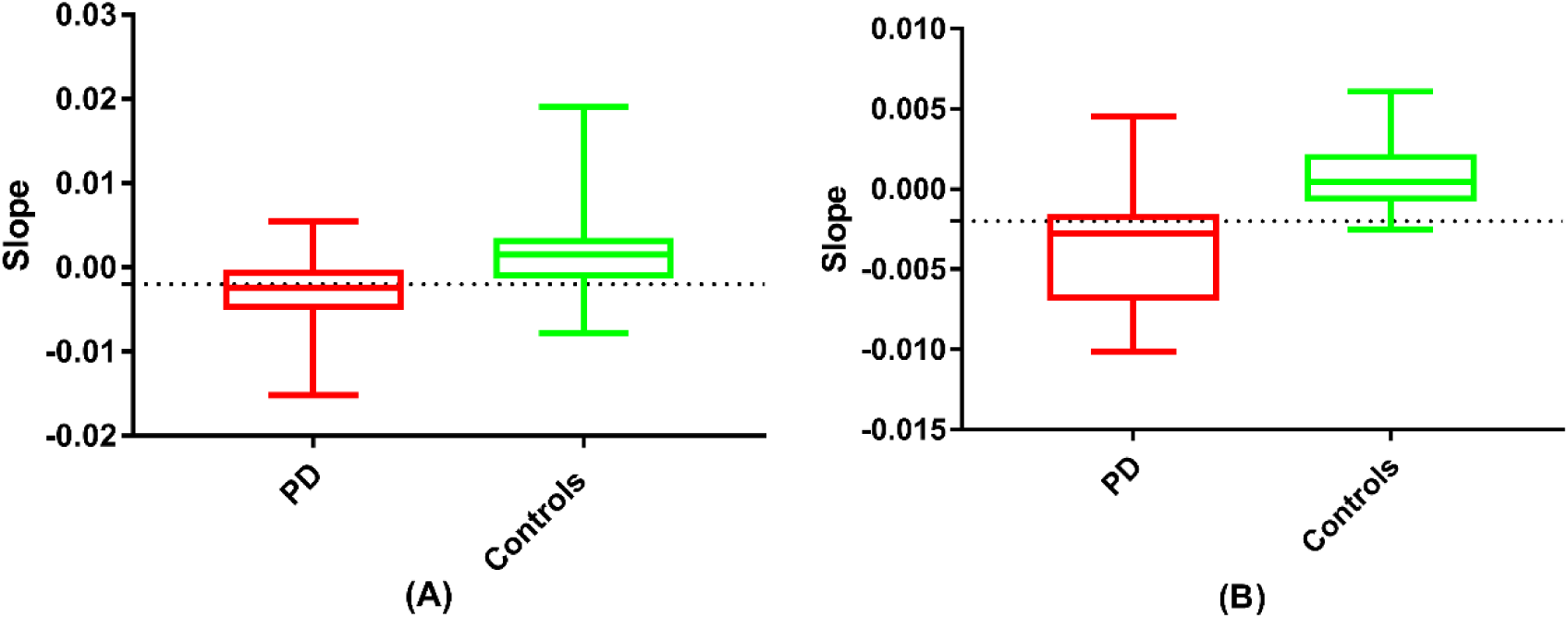
A box and whisker plot comparing: (A) PD patients in the ‘off state’ and most affected side (n = 60) and the non-dominant side for controls (n = 58) in the first experiment (Mann Whitney Test U = 841, p<0.001, two-tailed); (B) PD patients in the ‘off state’ and most affected side (n = 17) and controls non-dominant side (n = 17) in the third experiment (Mann Whitney Test U = 32, p<0.001, two-tailed). The cut-off is set at −0.002 which represents the 10^th^ centile cut-off for slopes in controls and the 50^th^ centile in PD patients with bradykinesia on clinical examination.

### Time-series analyses of dwell and travelling times for the sequence effect in PD

PD patients could not be differentiated from controls on the basis of a sequence effect of dwell and travelling times defined as ≥3 consecutive decrements in time series analyses of dwell and travelling times (see supplementary file, Tables S3 and S4).

### Differentiation between PD patients ‘off’ and ‘on’ medication

All BRAIN test parameters differentiated PD patients ‘off’ medication (n=61) and ‘on’ medication (see Table 2). Compared to PD patients’ ‘off state’ scores, patients ‘on’ medication had higher number of alternate taps (55.07±12.46 vs 49.11±11.34), lower average dwell times (110.4±34.22 vs 122.5±38.16 msec) and higher average tapping velocity (27.29±6.423 vs 24.17±5.563 cm/msec).

**Table 2.** Mean KS, Mean AT, Median IS, Median DS, Mean VS in PD patients (n = 61) and controls (n = 93) enrolled the second experiment. *P-values are reported for the differentiation between on and off states using Wilcoxon matched pairs signed rank test, **and between off state and controls using ROC curves. P-values reported were corrected for multiple testing using false discovery rate control (CI – Confidence Interval, IQR – Interquartile Range, PD – Parkinson’s disease, KS- kinesia Score, AT – akinesia Time, IS – incoordination Score, DS – dysmetria Score, VS – velocity Score)

| Parameter | PD 'off' | PD 'on' | p-value* | Controls | p-value** |
| --- | --- | --- | --- | --- | --- |
| Mean KS in taps [95% C.I] | 49.11<br>[46.5-51.6] | 55.07<br>[52.1-57.9] | <0.001 | 60.3<br>[57.6-63] | <0.001 |
| Mean AT in msec [95% C.I] | 122.5<br>[114.1-131] | 110.4<br>[102.9-118] | <0.001 | 112.1<br>[101.8-122.3] | 0.0008 |
| Median IS in msec<sup>2</sup> [IQR] | 67494<br>[11275-340854] | 167786<br>[33318-609816] | 0.0397 | 6758<br>[4030-16664] | <0.001 |
| Median DS in points [IQR] | 1.044<br>[1.018-1.136] | 1.097<br>[1.026-1.205] | 0.0003 | 1.044<br>[1.013-1.11] | 0.7636 |
| Mean VS in cm/sec [95% C.I] | 24.25<br>[22.99-25.5] | 27.29<br>[25.77-28.81] | <0.001 | 30.04<br>[28.31-31.76] | <0.001 |

### Differentiation between PD and controls

In the first and second experiments, KS, AT and IS scores differentiated between PD patients who were ‘off’ medication (n = 61) and healthy controls (n = 93). The same was observed with VS (n = 61) when scores in the ‘off state’ were compared to controls (n = 40) (supplementary material, Figure S3 A-D). DS did not convince in its ability to differentiate between the two groups (AUC = 0.52, p = 0.7636). IS offered the best discrimination between the two groups with sensitivities of 67%, 65% and 57% for specificities at 80%, 85% and 90% respectively. KS and VS were comparable offering sensitivities of 63%,59% and 26% and 60%,48% and 25% for specificities at 80%, 85% and 90%.

Similarly, in the third experiment, BRAIN test parameters differentiated between PD (n = 19) and controls (n = 19) consistently across the three platforms – keyboard, iPad and iPhone (except for AT parameter on the iPad, p = 0.088) (see Table 3). Area under the curve (AUC) values for Receiver Operating Characteristic (ROC) curve are summarised in supplementary material (Table S5).

**Table 3.** Mean KS, Median AT, IS, DS, DS1 and VS in PD patients (n = 19) and controls (n = 19) enrolled in the third experiment (CI – Confidence Interval, IQR – Interquartile Range, PD – Parkinson’s disease, KS- kinesia score, AT – akinesia time, IS – incoordination score, DS – dysmetria score, VS – Velocity Score).

| Platform | Mean KS (taps) (95% CI) |  |  | Median AT (msec) (IQR) |  |  | Median IS (msec<sup>2</sup>) (IQR) |  |  |
| --- | --- | --- | --- | --- | --- | --- | --- | --- | --- |
|  | <u>PD</u> | <u>Controls</u> | <u>p value</u> | <u>PD</u> | <u>Controls</u> | <u>p-value</u> | <u>PD</u> | <u>Controls</u> | <u>p value</u> |
| Standard keyboard | 51.47<br>(49.34,53.6) | 64.99<br>(62.53,67.45) | <0.001 | 107.5<br>(76.25,138.8) | 81<br>(62,106) | 0.0002 | 6585<br>(2870,20744) | 2366<br>(1328,4790) | <0.001 |
| iPad | 103.1<br>(96.82,109.5) | 120<br>(115.2,124.8) | <0.001 | 78.31<br>(65.7,91.24) | 73.47<br>(68.23,80.94) | 0.0879 | 1401<br>(746.7,2422) | 690.7<br>(483.5,1425) | <0.001 |
| iPhone | 100.6<br>(94.23,107.1) | 117.1<br>(112,122.2) | <0.001 | 88.14<br>(77.49,103.2) | 76.8<br>(68.75,88.11) | 0.0002 | 1304<br>(869.1,2863) | 1139<br>(494.3,2501) | 0.0371 |
|  | Median DS (points) (IQR) |  |  | Median DS1 (points) (IQR) |  |  | Mean VS (keys/sec) (95% C.I) |  |  |
|  | <u>PD</u> | <u>Controls</u> | <u>p value</u> | <u>PD</u> | <u>Controls</u> | <u>p-value</u> | <u>PD</u> | <u>Controls</u> | <u>p value</u> |
| Standard keyboard | 1.043<br>(1.003,1.124) | 1.021<br>(1,1.046) | 0.0023 | - | - | - | 16.88<br>(16,17.76) | 21.18<br>(20.41,21.95) | <0.001 |
| iPad | 1<br>(1,1.045) | 1 (1,1.008) | 0.0031 | 0.1241<br>(0.0763,0.1812) | 0.0843<br>(0.0652,0.1066) | <0.001 | - | - | - |
| iPhone | 1.027<br>(1.009,1.129) | 1 (1,1.028) | <0.001 | 0.1450<br>(0.122,0.1952) | 0.1007<br>(0.0834,0.1404) | <0.001 | - | - | - |

### Correlation with total motor UPDRS scores and sub-scores

KS and VS showed moderate inverse correlation with total motor scores of the MDS-UPDRS and sub-scores (pronation/supination, finger tapping, hand movements and upper limb rigidity) in both ‘on’ and ‘off’ states but other parameters lacked evidence of an association (see supplementary material Figure S4 A-D, Tables S6 and S7). Of the two parameters, VS showed marginally stronger correlation than KS.

### Test-retest reliability

ICC values with corresponding 95% confidence intervals are summarised in the supplementary material (Table S8). Using the keyboard version, all BRAIN tap parameters except for IS (poor reliability ICC = 0.141, p = 0.138), achieved good reliability (KS ICC = 0.881, DS = 0.808, VS = 0.883, p<0.001) and excellent reliability (AT ICC = 0.929, p<0.001). With the tablet device and smartphone, only KS (ICC = 0.836, p<0.001) and AT (ICC = 0.760, p<0.001) achieved good reliability.

## Discussion

Here, we report data using the BRAIN test, which address outstanding questions from earlier assessments [13,14]. We demonstrate a new measure for bradykinesia (sequence effect), using the VS which captures a decrement in repetitive movement, as opposed to the previous measures, which looked at speed of alternate tapping (KS) and dwell time (AT). The new VS parameter correlated best of all the five parameters with established parkinsonian signs and performed similarly to the KS and IS when differentiating patients from controls.

A key finding from this set of experiments was the ability for all BRAIN test parameters to differentiate between a patient ‘on’ and ‘off’ dopaminergic medication. This raises the possibility of using the BRAIN test to monitor motor fluctuations and to assist with therapeutic decision-making. Currently, decisions made clinically regarding efficacy of treatment depend on clinical examination, records of timing of medication and patient’s subjective reporting of symptoms and ability to perform activities of daily living on self-scoring diaries [21]. Identification of symptoms through history taking is affected by recall bias, together with the difficulty experienced by many patients in differentiating between normal, dyskinetic and bradykinetic states [22].

At the chosen cut-off slope of −0.002, the false positive rate was minimised to ~10% and the detection rate was ~60%. The reason for the sub-optimal detection rate in those with established PD may be due to the nature of alternate tapping and the fact that only proximal sequence effect can be detected in this setting (i.e. that which arises from movement at the shoulder/elbow). Adaptation of the test to better capture a distal sequence effect may be beneficial in a future iteration.

BRAIN tap test parameters correlated only approximately with MDS-UPDRS part III scores. This could be because BRAIN tap test parameters such as KS (proxy measure for total taps), AT (dwell time) and DS (proxy measure for accuracy) capture aspects of bradykinesia not tested with MDS-UPDRS sub-scores (finger tapping, hand movements, pronation/supination) which focus on rhythm, slowing and decrement in amplitude [2,23]. IS provided the best differentiation between PD patients and controls. However, this was non-specific as only KS and VS correlated with recognised parkinsonian signs. BRAIN tap test parameters have high ICC values (>0.75), offering reassurance about the reliability of test scores when the test may be used at home or without supervision.

We have also introduced a version of the tapping task to ‘smart’ device platforms such as the iPad (tablet) and iPhone (smartphone). With the exception of AT, BRAIN tap parameters offered better differentiation between PD patients and controls using standard keyboard when compared to smart devices (see supplementary material, Table S5). Considering the increasing availability of these technologies, this a further step towards portable domiciliary and clinic-based testing. The BRAIN test requires no specialised hardware to be purchased and can be accessed online free of charge requiring 20 seconds for practice session and 1 minute to perform the test (30 seconds each hand).

This study has several limitations. Data mining/exploratory testing has a higher chance of obtaining false positive results when compared to hypothesis driven testing. However, this was corrected by using a false discovery rate control for conducting statistical tests [19]. PD is a multi-system disease and motor impairment affecting the lower limb, dyskinesia, rigidity and tremor are not captured by the test. The BRAIN tap test requires hand-eye coordination and problems may arise in patients with visual problems or severe tremor.

The BRAIN tap test is a simple, sensitive, reliable test of upper limb motor function in PD. It is free to use and has been validated against the accepted gold standard MDS-UPDRS part III rating scale. It can differentiate between ‘on’ and ‘off’ states in individual patients and can quantify a sequence effect using decrement in the VS score, making it a useful adjunctive outcome measure for clinical practice and in clinical trials.

## Acknowledgements

The work presented here had no specific funding but the Preventive Neurology Unit is funded by the Barts Charity, the Exenatide PD study was funded by the Michael J Fox Foundation, and the Cure Parkinson’s Trust. Dr. Noyce was funded by Parkinson’s UK at the time that the data were collected and M Burrows by the Reta Lila Weston Trust.

## Ethical standards

Studies have been approved by the appropriate ethics committee and have, therefore, been performed in accordance with the ethical standards laid down in the 1964 Declaration of Helsinki and its later amendments. All persons gave their informed consent prior to their inclusion in the study.

